## Supplemental Information for "Regulation of Chromatin Modifications through Coordination of Nucleus Size and Epithelial Cell Morphology Heterogeneity"

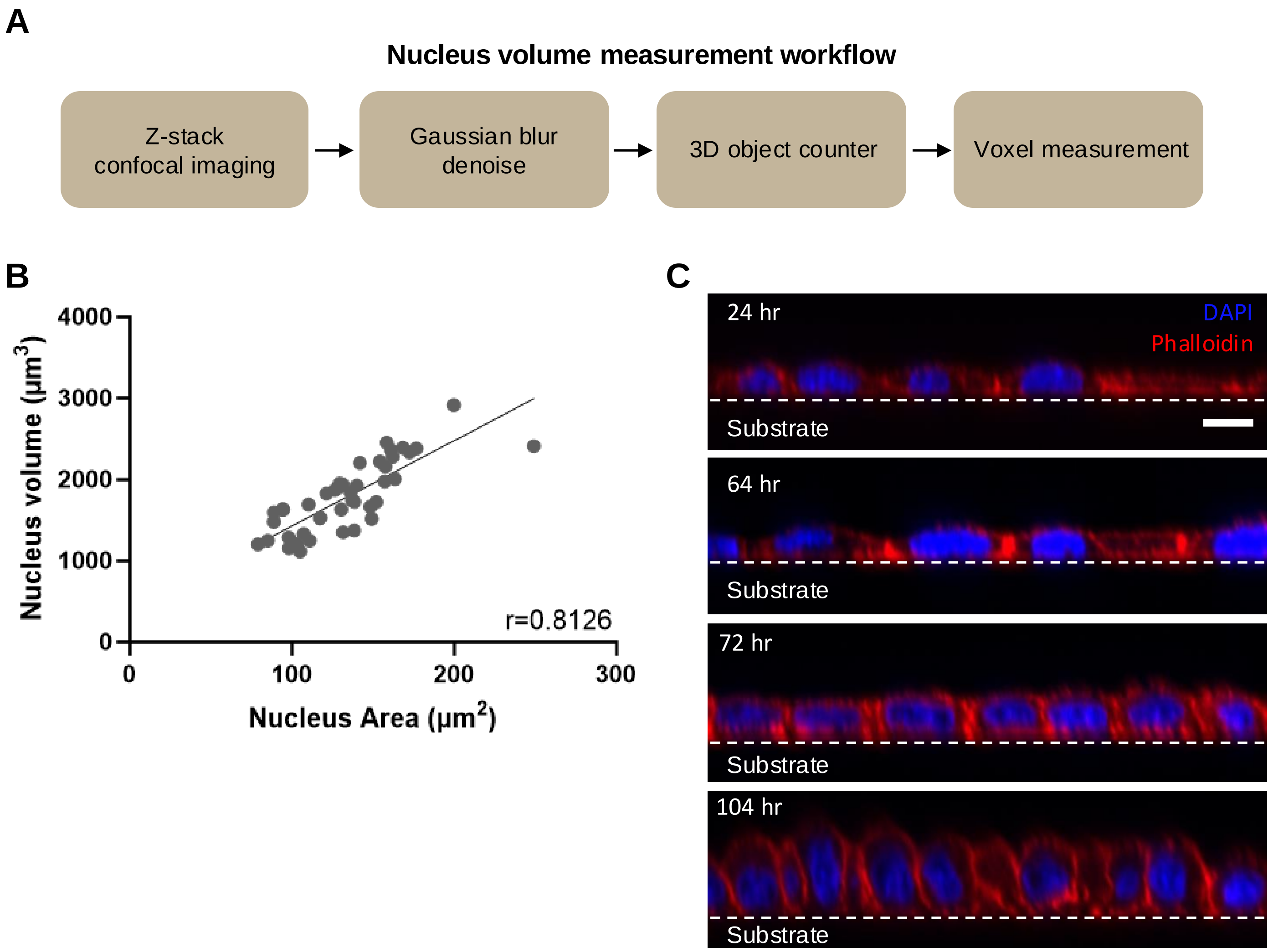


**Fig. S1. Area can be used to approximate volume in MDCK cells.** (A) Analysis pipeline overview for volume measurement acquisition. First, z-stack imaging is performed using a fine 0.6 μm step size. Next, the image stack is imported into ImageJ and duplicated. A 3D Gaussian blur is then applied the duplicate stack using $\sigma_{x,y,z}=2$. This blurred duplicate is then used to set the masking threshold using the 3D objects counter analysis feature. Measurements are performed on the unfiltered z-stack. After the 3D objects counter analysis is performed, the 3D voxel measurement is output for each nucleus. (B) High correlation between nucleus volume and nucleus area in confluent MDCK cells suggests that area can be used in place of volume to characterize cell morphological behavior. (C) Orthogonal view of MDCK cells throughout crowding illustrating sample flatness. Time stamp refers to time post seeding at 30k cells/cm$^2$. Scale bar 10 μm.

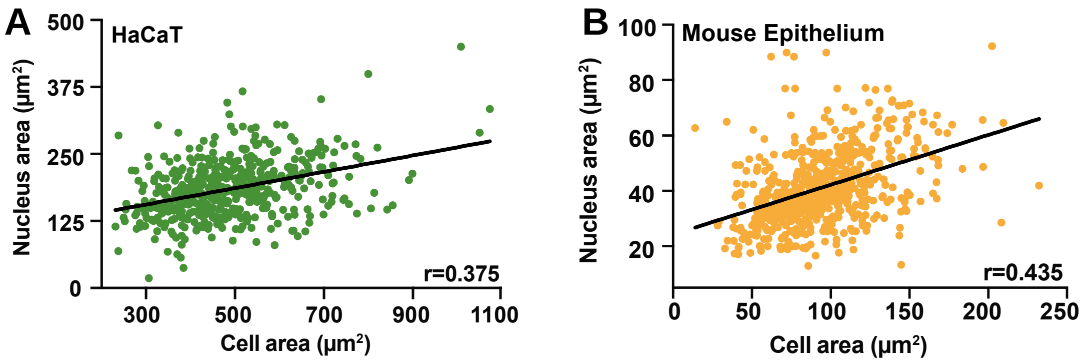


**Fig. S2. Nucleus-to-cell (NC) area correlations are demonstrated in HaCaT and mouse epithelium, akin to MDCK results.** NC area correlation exhibited in (A) confluent HaCaT cells and (B) E12.5 mouse arm epithelium.

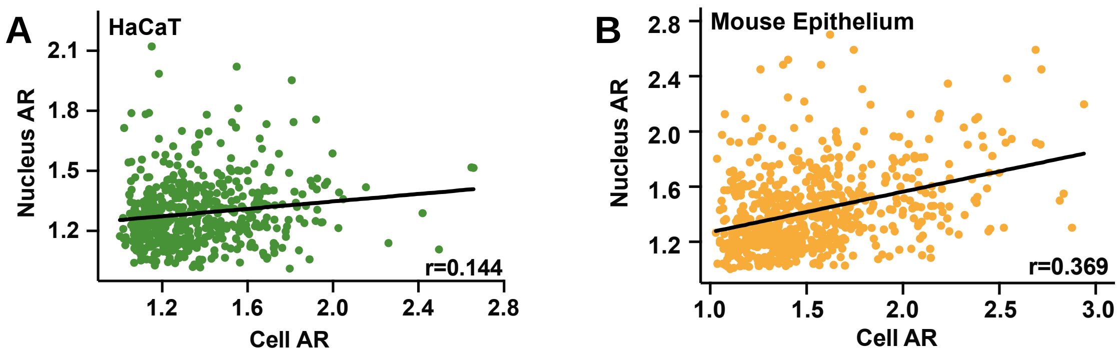


**Fig. S3. Nucleus-to-cell (NC) aspect ratio (AR) correlations are demonstrated in HaCaT and mouse epithelium, akin to MDCK results.** NC AR correlation exhibited in (A) confluent HaCaT cells and (B) E12.5 mouse arm epithelium.

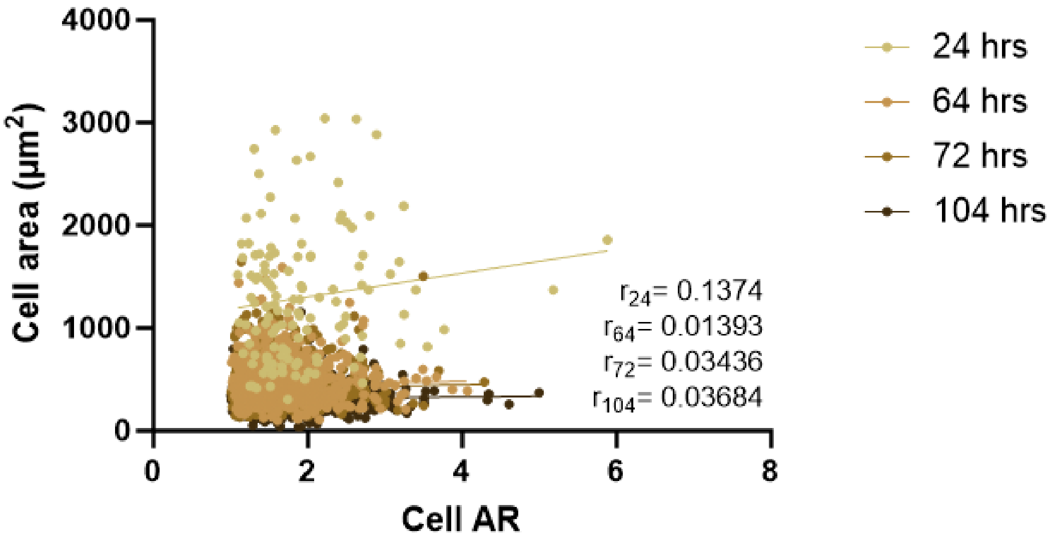


**Fig. S4. Cell area correlation with cell aspect ratio (AR) in MDCK cells.** Negligible correlation between cell area and AR is observed at all investigated time points, suggesting that these two morphological features are independent and regulated by distinctive mechanisms.

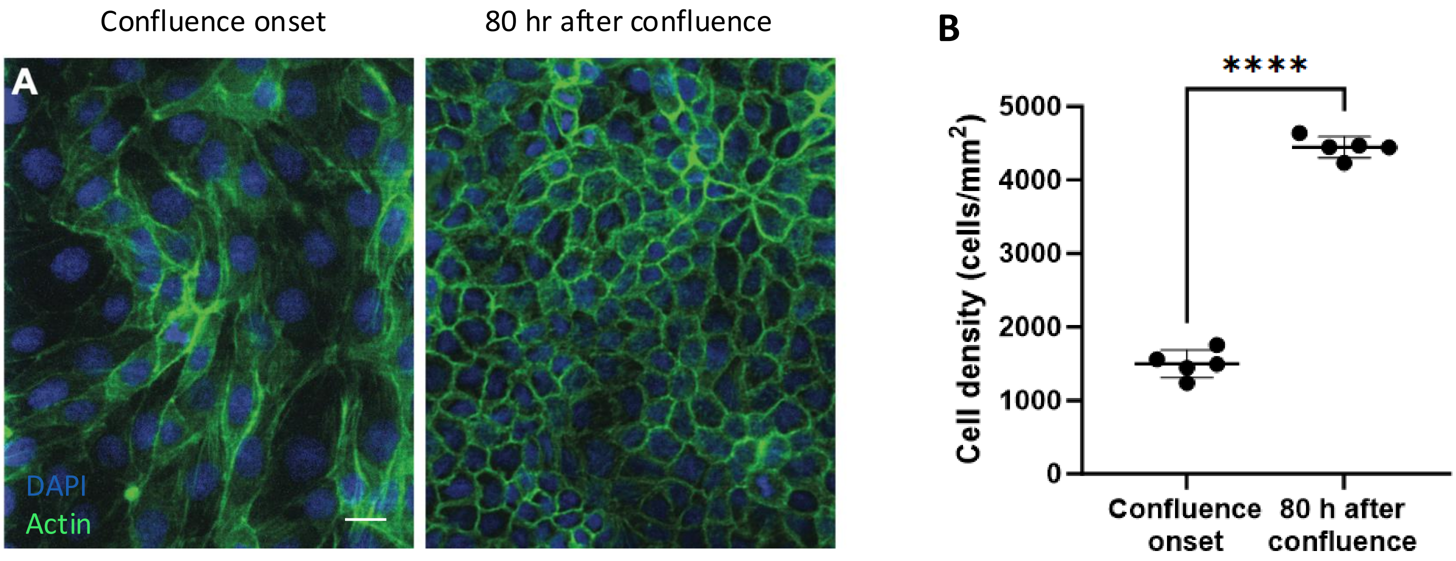


**Fig. S5. Cell division is required to induce cell crowding, and thus required for cell size reduction during crowding.** (A) Confluent MDCK monolayer (left) has less cells and larger cells compared to a crowded monolayer (right). Scale bar 20 µm. (B) Cell density approximately triples 80 hours after reaching confluence, suggesting that cell division and cell area reduction can be coupled. Error bars indicate the standard deviation.

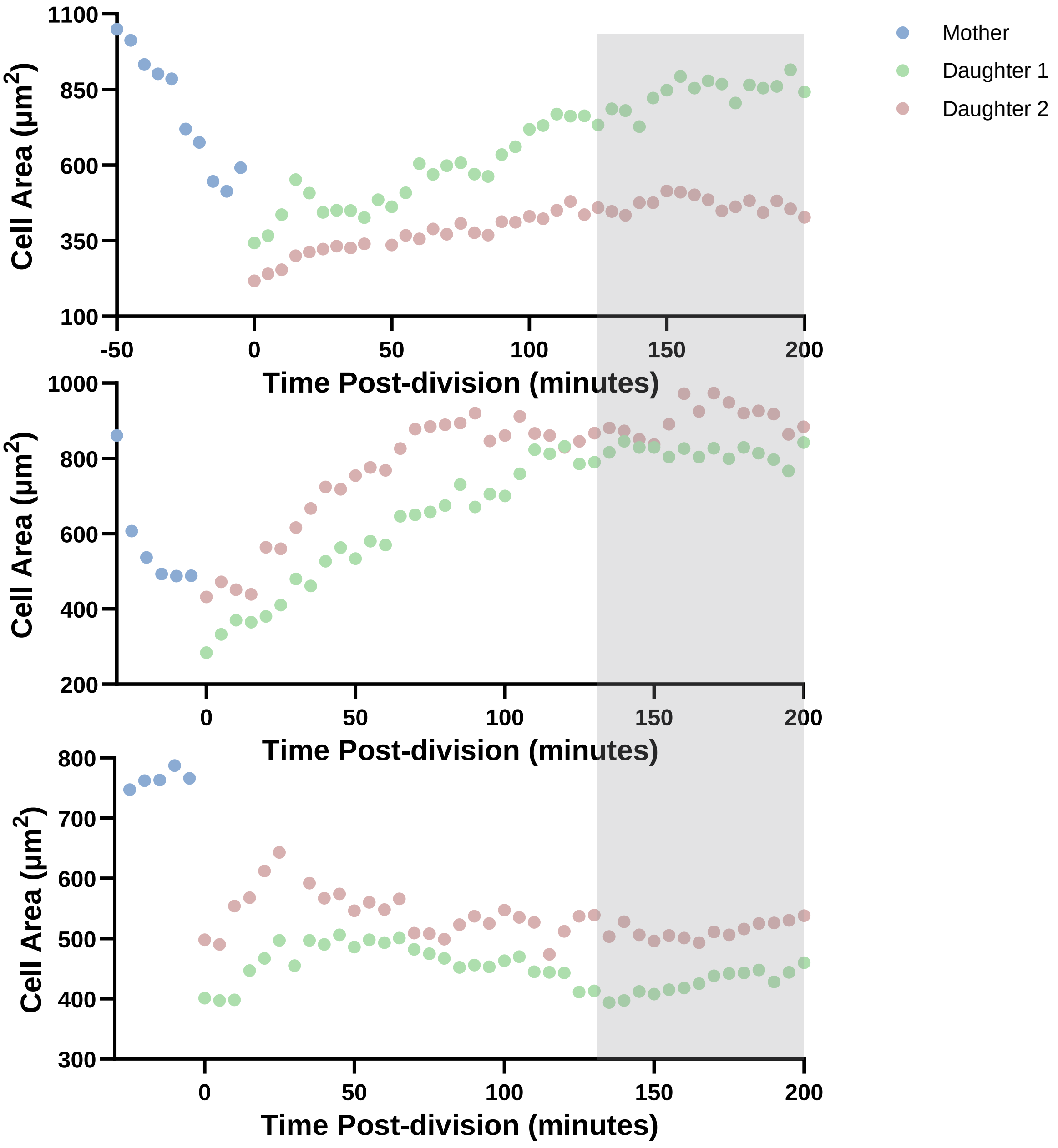


**Fig. S6. Cell growth rate plateaus approximately 2 hours post division.** Three representative examples of cell growth trajectories illustrate that the cell growth phase lasts approximately 2 hours after division for most of the analyzed cell pairs. Gray shaded band indicates growth plateau regime.

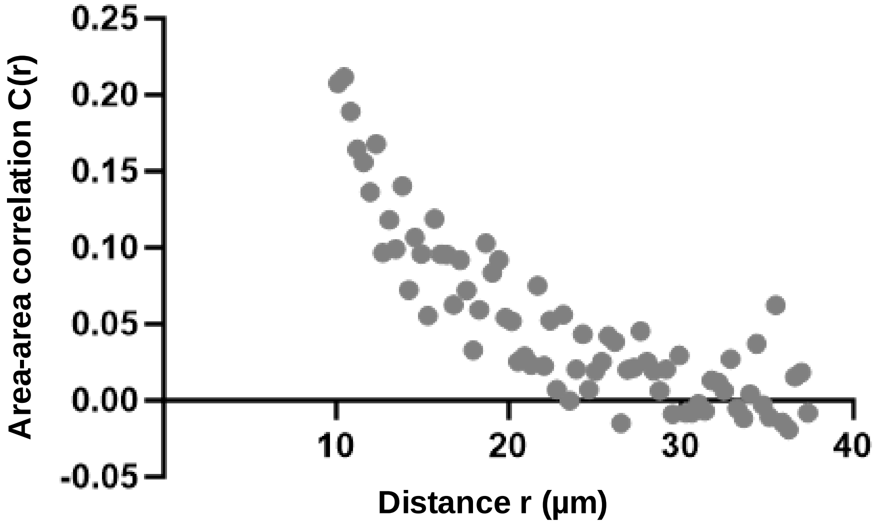


**Fig. S7. Area-area autocorrelation function of crowded MDCK cells reveals essentially no correlation between neighboring cells.** For distances greater than the diameter of a crowded MDCK cell (~10-15 μm), no significant correlation is observed, suggesting that there is no genetic memory or inheritance of final cell size. The correlation $C(r)$ is calculated using $C(r) = \sum_{i,j} \frac{(A(r_i)-\langle A \rangle)(A(r_j)-\langle A \rangle)}{\langle A^2 \rangle - \langle A \rangle^2}$, where $i, r,$ A and$\langle X \rangle$ denote the cell index, distance, area, and arithmetic mean, respectively.

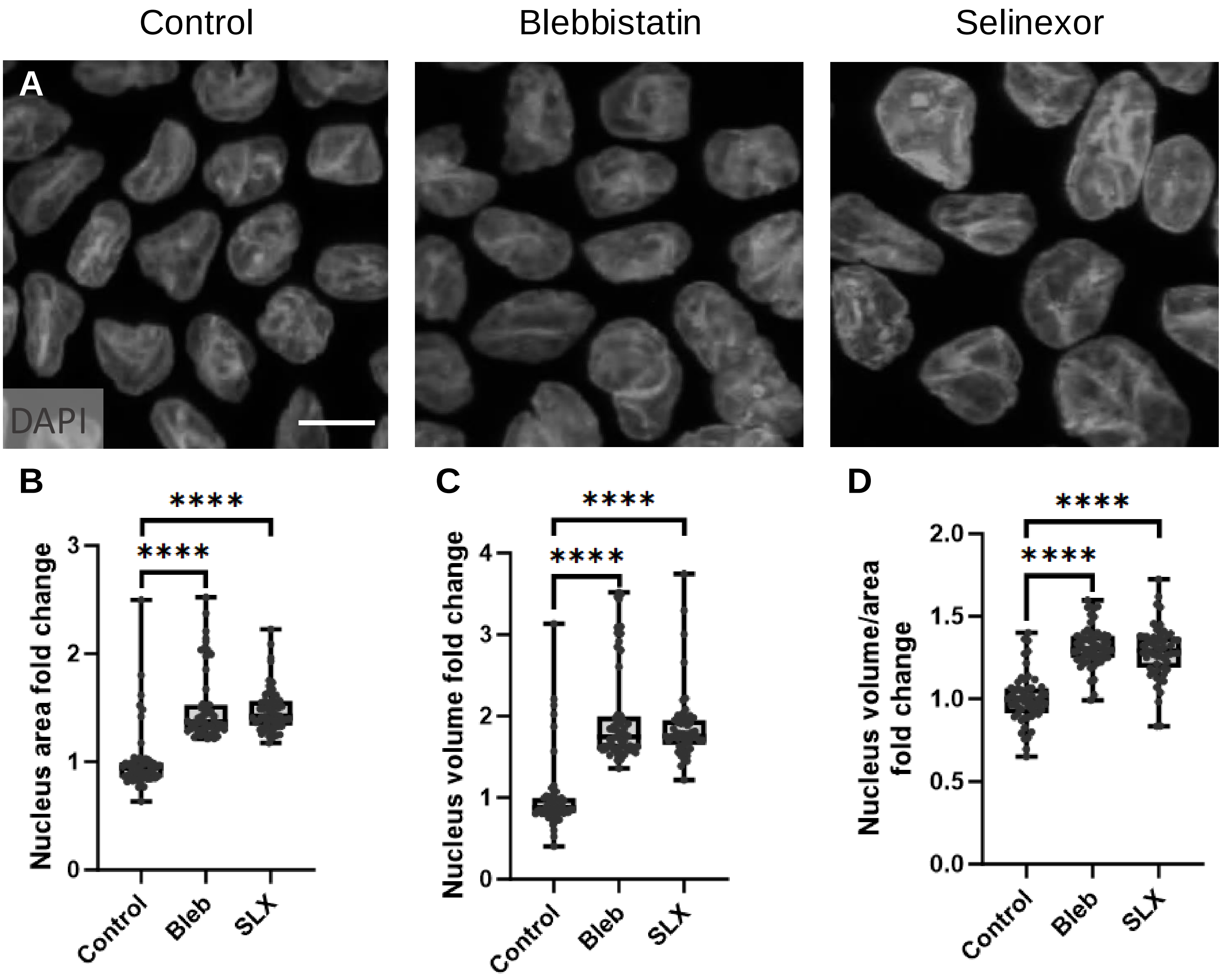


**Fig. S8. Nucleus volume increases in response to blebbistatin (bleb) and Selinexor (SLX) treatments.** (A) 3D projected confocal immunofluorescence images of untreated, bleb, and SLX treated nuclei, illustrating size differences. Scale bar 10 µm. (B) Nucleus area quantification demonstrates a ~1.4 fold increase in nucleus area in treated groups. (C) Such a size increase is also reflected in nucleus volume, which was measured using the Imaris volume measurement function. Specifically, we found that bleb- and SLX-treated cells exhibit a ~1.75 fold increase in nucleus volume. (D) Nucleus volume/area is used to estimate nucleus height, in which a slight (~1.25 fold) increase was observed in both treated groups. Treatment concentration was 10 µM for both bleb and SLX.

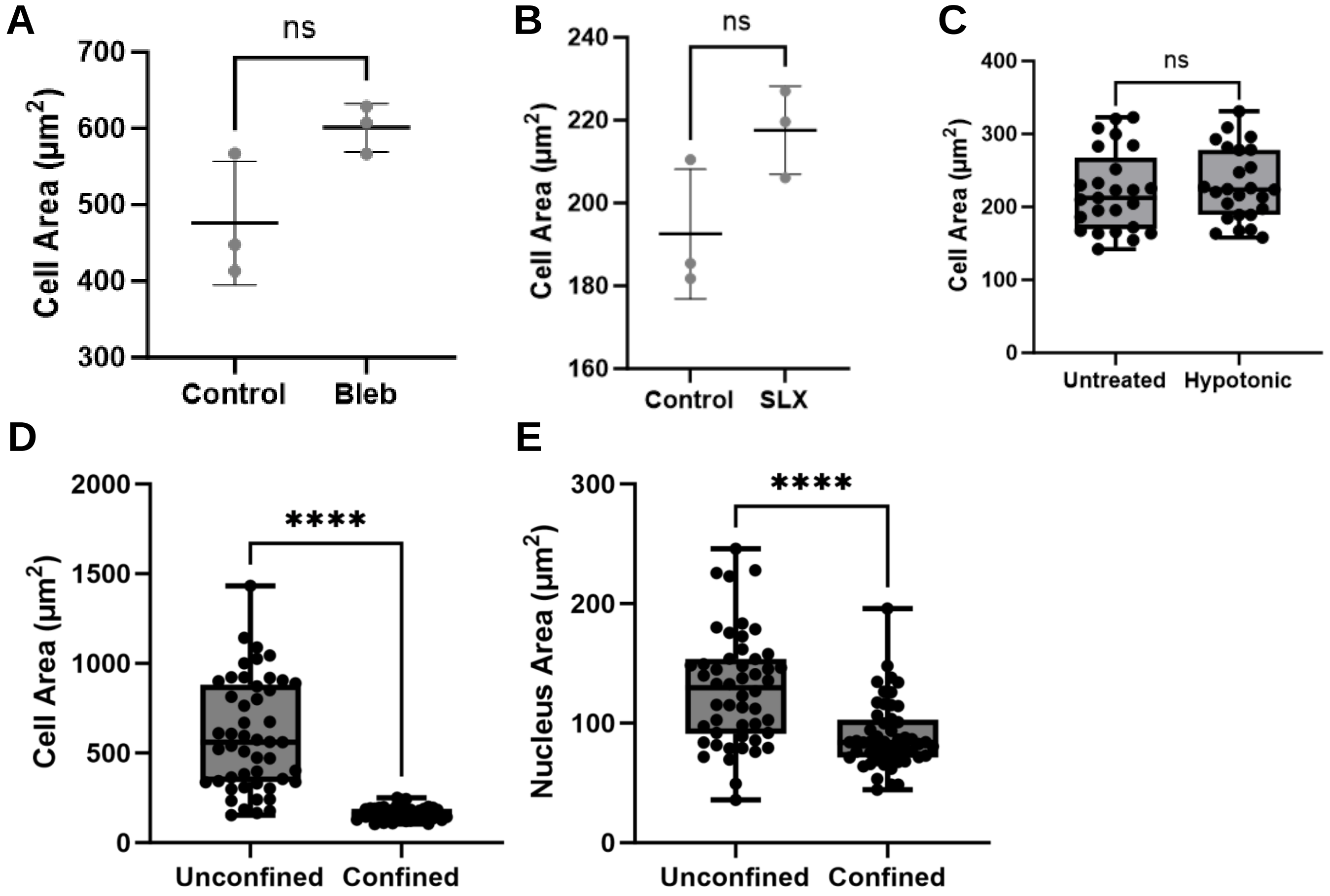


**Fig S9. Cell area remains unchanged by blebbistatin, Selinexor, and hypotonic treatment while confinement reduces both nucleus and cell size.** (A) Quantification of cell area for control and blebbistatin treated cells**.** (B) Same as (A) but with selinexor treated cells. (C) Same as (A) but with hypotonic shocked cells. (D) Quantification of cell area for control and 10 µm-confined cells**.** (E) Same as (D) but for nucleus area. Error bars indicate the standard deviation. ”ns” refers to a p-value ≥ 0.05.

A

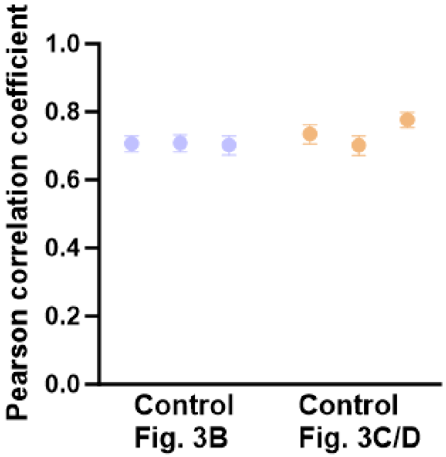


B

| Z-test values | 3b-Rep 1 | 3b-Rep 2 | 3b-Rep 3 | 3c/d-Rep 1 | 3c/d-Rep 2 | 3c/d-Rep 3 |
|---|---|---|---|---|---|---|
| 3b-Rep 1 | | -0.093 | 0.194 | -1.497 | 0.268 | -4.191 |
| 3b-Rep 2 | | | 0.275 | -1.379 | 0.345 | -3.992 |
| 3b-Rep 3 | | | | -1.588 | 0.075 | -4.131 |
| 3c/d-Rep 1 | | | | | 1.620 | -2.304 |
| 3c/d-Rep 2 | | | | | | -4.098 |
| 3c/d-Rep 3 | | | | | | |

C

| p-values | 3b-Rep 1 | 3b-Rep 2 | 3b-Rep 3 | 3c/d-Rep 1 | 3c/d-Rep 2 | 3c/d-Rep 3 |
|---|---|---|---|---|---|---|
| 3b-Rep 1 | | 0.92608 | 0.84598 | 0.13439 | 0.78853 | 0.00003 |
| 3b-Rep 2 | | | 0.78332 | 0.16792 | 0.73038 | 0.00007 |
| 3b-Rep 3 | | | | 0.11239 | 0.93986 | 0.00004 |
| 3c/d-Rep 1 | | | | | 0.10533 | 0.02124 |
| 3c/d-Rep 2 | | | | | | 0.00004 |
| 3c/d-Rep 3 | | | | | | |

D

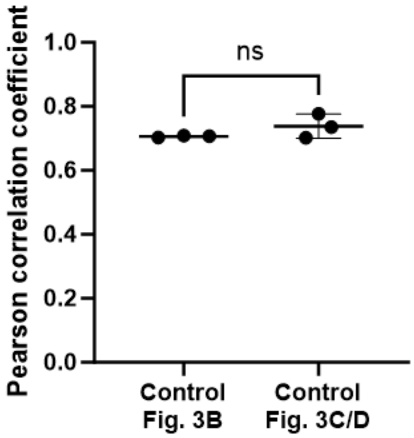


**Fig S10. Control nucleus area - cell area Pearson correlation coefficient analysis.** (a) Pearson correlation coefficients for the six biological replicates across two independent control experiments shown in Figs. 3B and 3C/D. The coefficients exhibit similar values, with variations generally within the confidence intervals indicated by the error bars. (b) Summary of statistical significance analysis using Fisher's Z-transformation and Z-test, comparing correlations between replicates. Z-test values greater than 1.96 or less than -1.96 indicate significant differences (yellow shaded). (c) Converted p-values from the Z-test, based on a two-tailed probability using the standard normal distribution. The analysis reveals a significant difference in the final replicate, while the other datasets show no significant differences, suggesting overall consistency in the Pearson correlation coefficients across the six biological replicates. (d) Additionally, to assess batch-to-batch variation in the Pearson correlation coefficient measurements, we calculated the p-value between the two independent control experiments. The analysis revealed no significant difference between the two experiments, as shown in the final figure below. Error bars indicate the standard deviation. "ns" denotes a p-value ≥ 0.05.

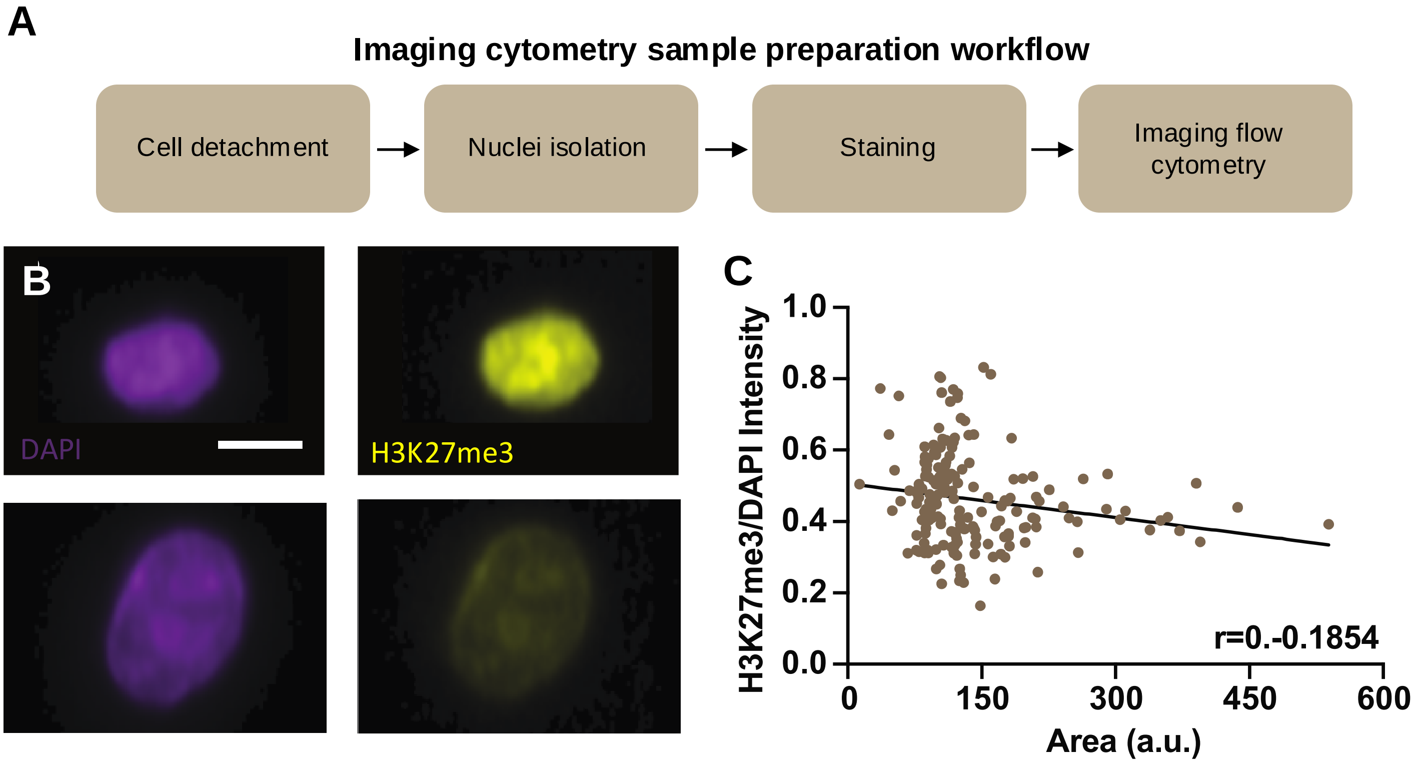


**Fig. S11. Anti-correlation between histone mark correlation with nucleus area in MDCK cells analyzed using imaging flow cytometry (Image Stream).** (A) Experiment schematic for preparing samples for imaging flow cytometry analysis. First, cells were detached from the substrate using trypsin before being pelleted by centrifugation. The supernatant was aspirated, and cells were resuspended in a cell lysis buffer containing 0.1% NP40, 0.01% digitonin, and 0.1% Tween20. After a 5-minute incubation on ice, the lysis buffer was quenched using a resuspension buffer containing ultra pure water, 1M Tris0HCL, 5M NaCl, 1M $MgCl_2$, and 0.1% Tween20. Resulting nuclei were then centrifuged to remove supernatant and resuspended in formalin for 10 minutes to perform fixation. Nuclei were then centrifuged to remove the formalin and washed twice using PBS and centrifugation. After the last wash, nuclei were stained for DAPI and H3K27me3 as described in the Materials and Methods section. These samples were then used for imaging flow cytometry analysis. (B) Representative images from imaging flow cytometry analysis demonstrating small and large nuclei express more and less H3K27me3, respectively. (C) Analysis of imaging flow cytometry data quantifying anti-correlation between normalized H3K27me3 intensity and nucleus size.

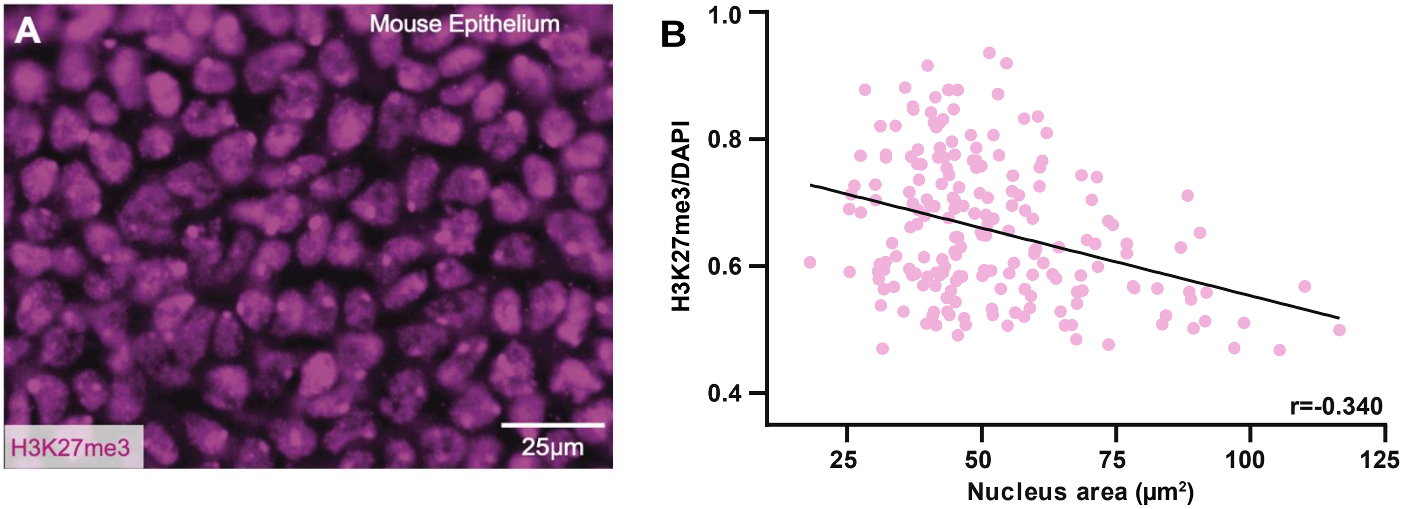


**Fig. S12. H3K27me3 intensity analysis in E11.5 mouse epithelium demonstrates that H3K27me3 is anti-correlated with nucleus size, in agreement with the MDCK result and other mouse epithelium results at E12.** (A) Representative image of mouse arm epithelium stained with H3K27me3. (B) Anti-correlation between normalized H3K27me3 intensity and nucleus size.

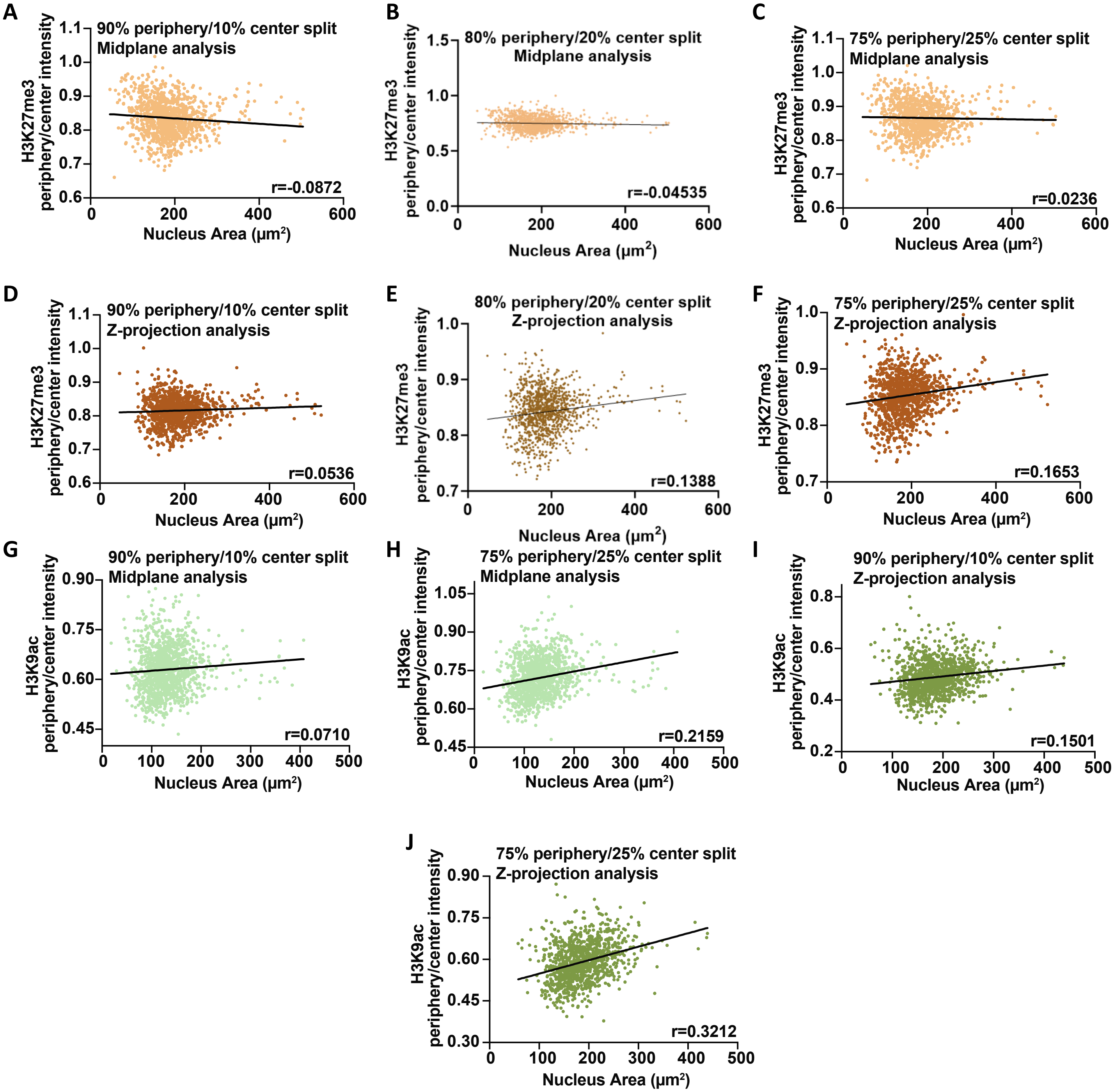


**Fig. S13. H3K27me3 and H3K9ac spatial distribution analysis in MDCK cells**. H3K27me3 localization within the nucleus was examined using a single midplane image using a split ratio in which either (A) the inner 90% of the nucleus was considered the center and the outer 10% was considered the periphery, (B) the inner 80% of the nucleus was considered the center and the outer 25% was considered the periphery, or (C) the inner 75% of the nucleus was considered the center and the outer 25% was considered the periphery. (D) Same analysis as (A) using a maximal z-projection instead of a single midplane slice. (E) Same analysis as (B) using a maximal z-projection instead of a single midplane slice. (F) Same analysis as (C) using a maximal z-projection instead of a single midplane slice. (G-J) Same analysis as A, C, D, and F, respectively, investigating H3K9ac instead of H3K27me3.

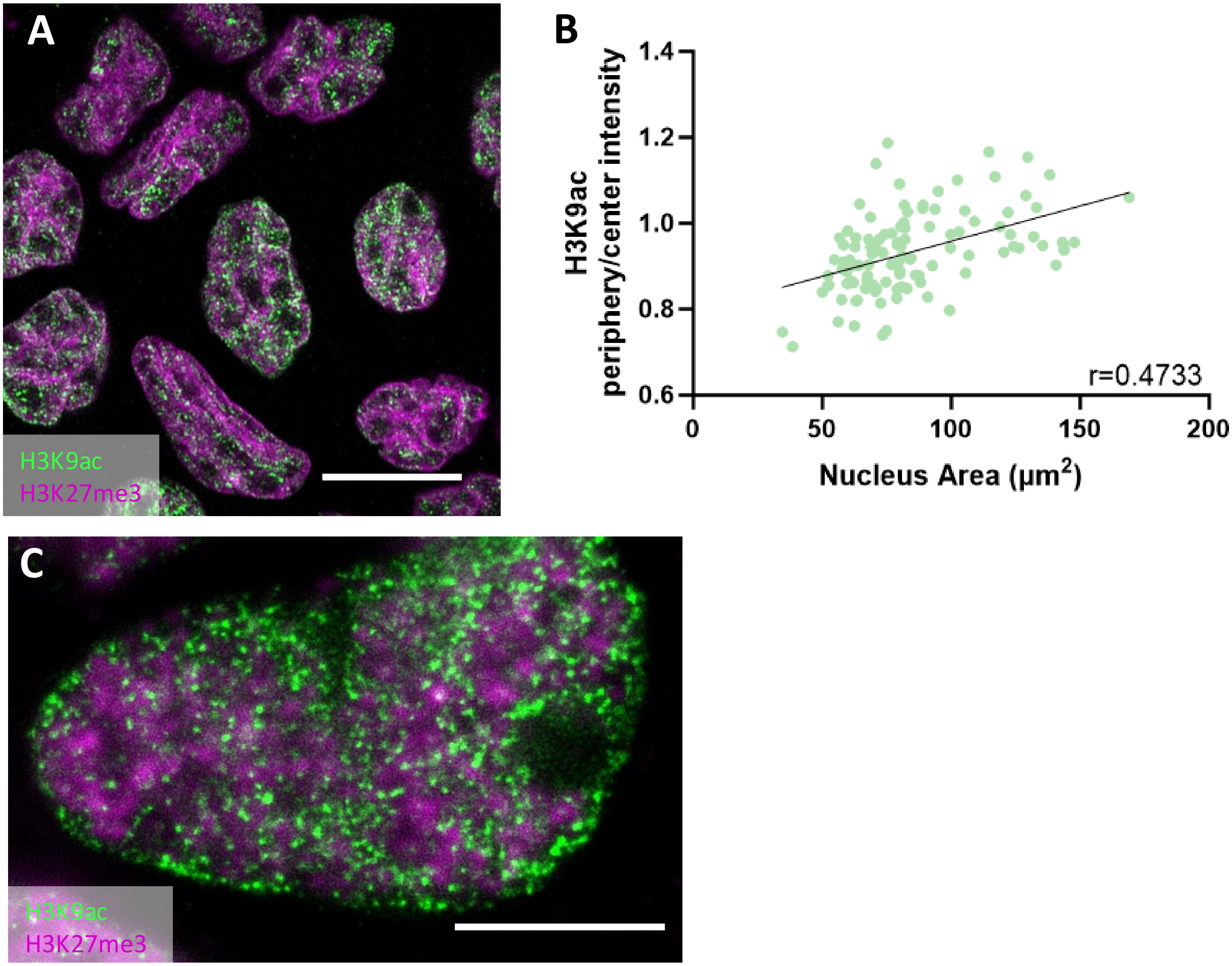


**Fig. S14. Super-resolution analysis of H3K27me3 and H3K9ac in crowded MDCK cells reproduces correlation between H3K9ac localization and nucleus area obtained from confocal microscopy.** (A) Airyscan images of MDCK nuclei illustrates distinct localization of active H3K9ac mark and repressive H3K27me3 mark. H3K9ac is localized as puncta within the nuclei while H3K27me3 localized around nuclear folds and edges. Scale bar 10 μm. (B) Radial analysis of H3K9ac using airyscan images. A positive correlation between H3K9ac periphery/center intensity and nucleus area reproduces confocal result shown in figure 4. n = 114 nuclei. p-value <0.0001. Confidence interval = [0.3169 to 0.6046]. (C) STED image of H3K27me3 and H3K9ac stained nucleus demonstrates histone mark localization consistent with airyscan and confocal results. Scale bar 5 μm.

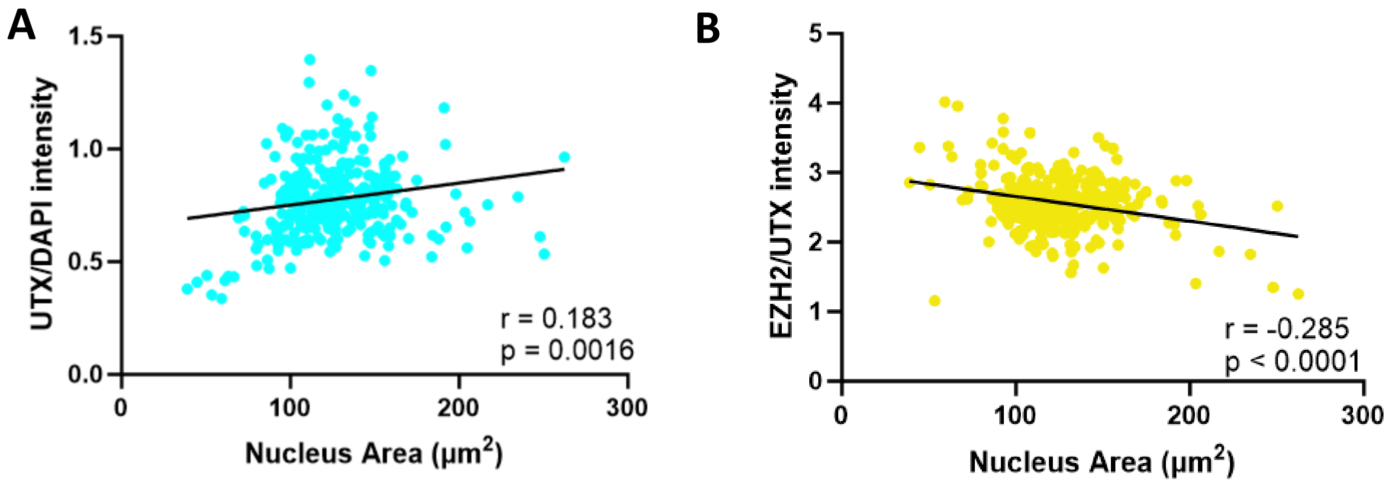


**Fig. S15. Histone demethylation UTX is correlated with nucleus size.** (A) UTX/DAPI intensity is positively correlated with Nucleus size. (B) EZH2/UTX intensity is anti-correlated with nucleus size.

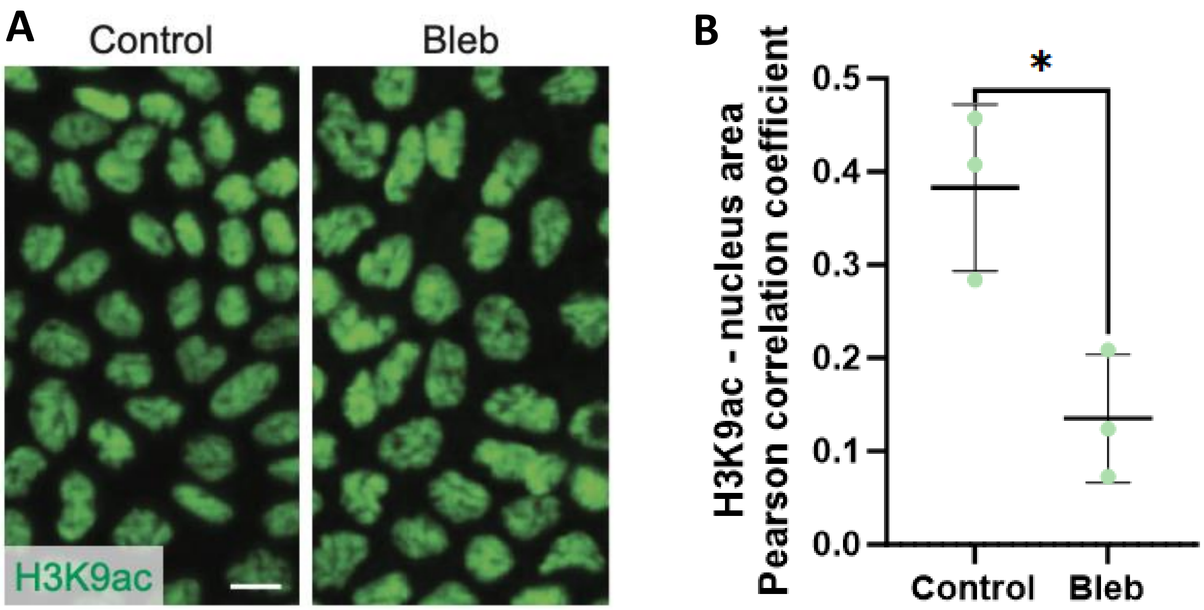


**Fig. S16. Myosin II is required for maintaining the positive correlation between nucleus area and H3K9ac level.** (A) Image of control (left) and Bleb-treated (right) MDCK cells stained with H3K9ac. Scale bar = 10 μm. (B) Pearson correlation coefficient between H3K9ac/DAPI intensity and nucleus area for control and Bleb-treated cells. Error bars indicate the standard deviation. * refers to p-value < 0.05.

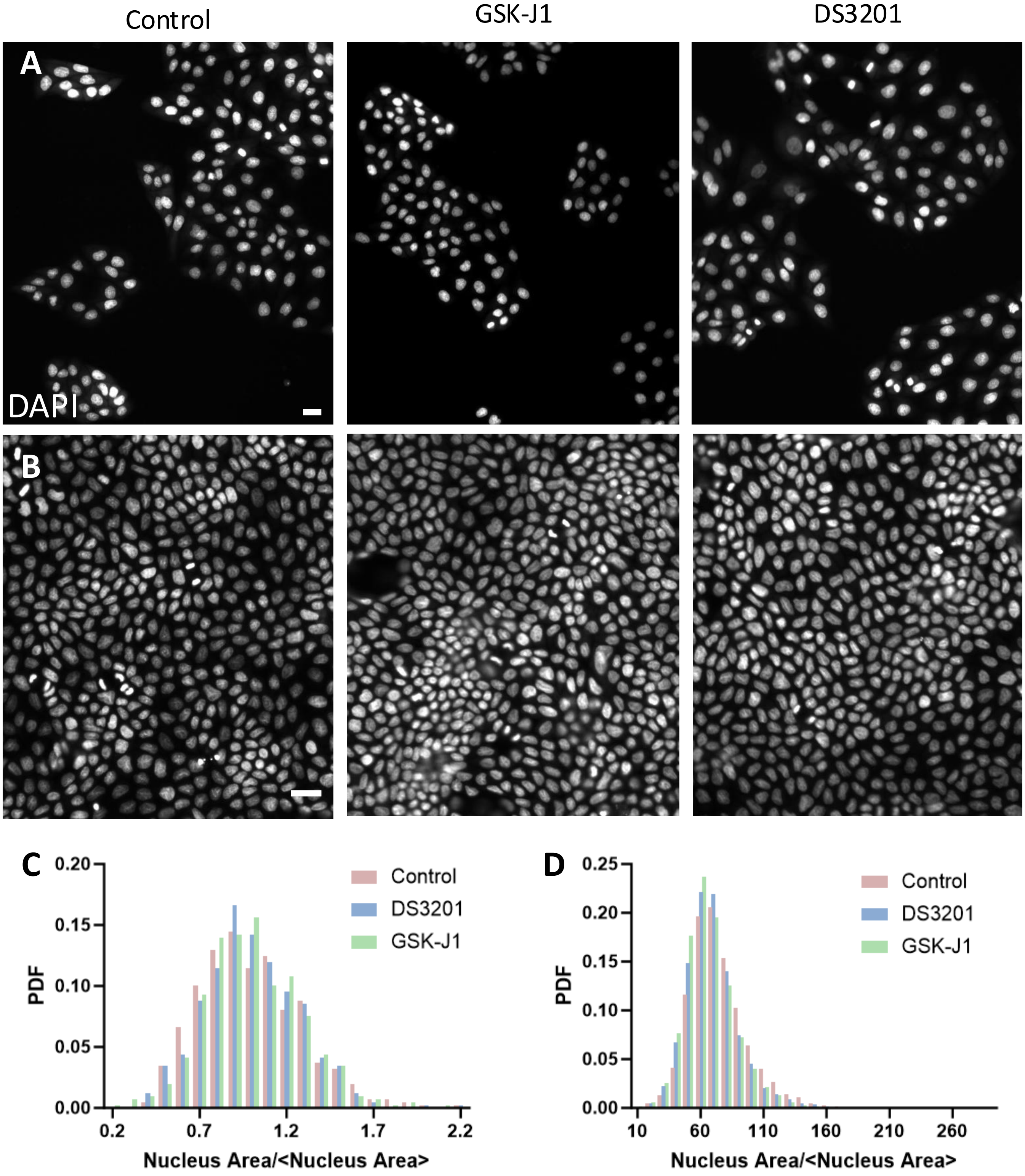


**Fig. S17. Nuclear size heterogeneity in subconfluent and crowded cells.** (A) Subconfluent MDCK nuclei in control (left), GSK-J1 (middle) and DS3201 (right) treated cells. (B) Crowded MDCK nuclei in control (left), GSK-J1 (middle) and DS3201 (right) treated cells. (C) Probability density function (PDF) of subconfluent MDCK nucleus area normalized to the mean nucleus area in control , GSK-J1 and DS3201 treated cells. (D) Same as (C) for crowded nuclei. Scale bar = 25 μm.

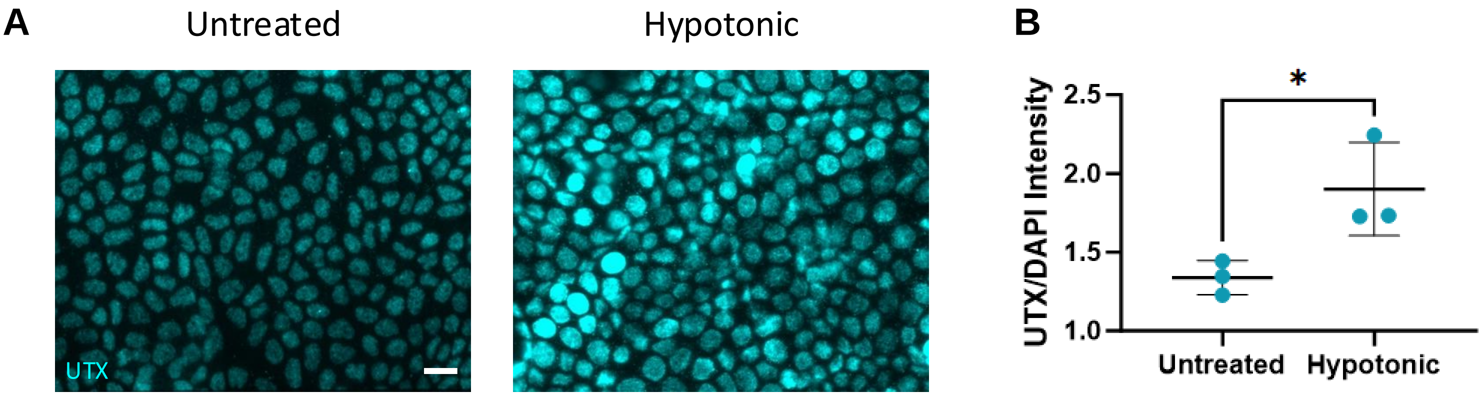


**Fig. S18. UTX transport is influenced by nucleus size.** (A) Fluorescent images of untreated cells (left) and cells after a 10-minute hypotonic shock treatment (right). (B) UTX intensity normalized to DAPI in untreated and hypotonic shocked cells. Error bars indicate the standard deviation. ”ns”, *, and ***, refer to p-values ≥ 0.05, < 0.05, and < 0.001, respectively.

| Figure | Pearson Correlation | Spearman Correlation | p-value | FDR | 95% Confidence Interval |
|---|---|---|---|---|---|
| Fig. 1D MDCK nucleus area vs. cell area (24 hr) | 0.6365 | 0.766 | <0.0001 | $<10^{-5}$ | [0.518, 0.731] |
| Fig. 1D MDCK nucleus area vs. cell area (64 hr) | 0.8326 | 0.838 | <0.0001 | $<10^{-5}$ | [0.810, 0.853] |
| Fig. 1D MDCK nucleus area vs. cell area (24 hr) | 0.7980 | 0.7506 | <0.0001 | $<10^{-5}$ | [0.771, 0.822] |
| Fig. 1D MDCK nucleus area vs. cell area (104 hr) | 0.7007 | 0.7066 | <0.0001 | $<10^{-5}$ | [0.668, 0.731] |
| Fig. 1H MDCK nucleus AR vs. cell AR (24 hr) | 0.3031 | 0.2359 | <0.0001 | $<10^{-5}$ | [0.134, 0.455] |
| Fig. 1H MDCK nucleus AR vs. cell AR (64 hr) | 0.4244 | 0.4047 | <0.0001 | $<10^{-5}$ | [0.366, 0.479] |
| Fig. 1H MDCK nucleus AR vs. cell AR (24 hr) | 0.4455 | 0.3858 | <0.0001 | $<10^{-5}$ | [0.388, 0.499] |
| Fig. 1H MDCK nucleus AR vs. cell AR (104 hr) | 0.5749 | 0.5272 | 0.0075 | $<10^{-5}$ | [0.533, 0.614] |
| Fig. 2D Cell area 6 hr vs. cell area 0 hr | 0.568 | 0.6087 | <0.0001 | $<10^{-5}$ | [0.431, 0.679] |
| Fig. 2E Nucleus area 6 hr vs. nucleus area 0 hr | 0.586 | 0.5711 | <0.0001 | $<10^{-5}$ | [0.452, 0.694] |
| Fig. 2H Nucleus area vs. cell area (0 hr) | 0.727 | 0.6985 | <0.0001 | $<10^{-5}$ | [0.628, 0.803] |
| Fig. 2I Nucleus area vs. cell area (6 hr) | 0.811 | 0.8444 | <0.0001 | $<10^{-5}$ | [0.738, 0.865] |
| Fig. 4B MDCK H3K27me3/DAPI vs. nucleus area | -0.326 | -0.3280 | <0.0001 | $<10^{-5}$ | [-0.447, -0.194] |
| Fig. 4D Mouse ectoderm H3K27me3/DAPI vs. nucleus area | -0.340 | -30.19 | <0.0001 | $<10^{-5}$ | [-0.459, -0.209] |
| Fig. 4F MDCK H3K9ac/DAPI vs. nucleus area | 0.239 | 0.2969 | <0.0001 | $<10^{-5}$ | [0.183, 0.293] |
| Fig. 4H Mouse ectoderm H3K9ac/DAPI vs. nucleus area | 0.171 | 0.1674 | <0.0001 | $<10^{-5}$ | [0.0988, 0.241] |
| Fig. 4K Mouse ectoderm H3K9ac P/C intensity vs. nucleus area | 0.185 | 0.1234 | <0.0001 | $<10^{-5}$ | [0.131, 0.238] |

**Table S1. Summary of correlative analyses.** To quantify correlation, both the Pearson (column 2) and Spearman (column 3) correlation coefficient were measured. To quantify statistical significance the p-value (column 4), false discovery rate (FDR) (column 5), and 95% confidence interval (column 6) were computed.
